# Beyond venomous fangs: Uloboridae spiders have lost their venom but not their toxicity

**DOI:** 10.1101/2023.06.26.546488

**Authors:** Xiaojing Peng, Ludwig Dersch, Josephine Dresler, Tim Lüddecke, Tim Dederichs, Peter Michalik, Steve Peigneur, Jan Tytgat, Afrah Hassan, Antonio Mucciolo, Marc Robinson-Rechavi, Giulia Zancolli

## Abstract

Venom, one of nature’s most potent secretions, has played a crucial role in the evolutionary success of many animal groups, including spiders. However, Uloboridae spiders appear to lack venom and capture their prey, unlike venomous spiders, by extensive silk-wrapping and regurgitation of digestive fluids onto the entire prey. A prevailing hypothesis posits that toxins may have been reallocated from the venom to alternative secretions, like silk or digestive fluids. Yet, whether uloborids have retained venom toxins and the mechanisms underlying prey immobilisation remain unresolved. Here, we employed a multi-disciplinary approach to assess the absence of venom glands in *Uluborus plumipes*, toxin gene expression and toxicity of digestive proteins. Our findings confirm that *U. plumipes* lacks a venom apparatus, while neurotoxin-like transcripts were highly expressed in the digestive system. Midgut gland extract had comparable toxicity levels to that of the venomous *Parasteatoda tepidariorum*. However, no inhibitory effects on sodium nor potassium channels were observed, indicating a different toxic mechanism. These findings support the hypothesis that Uloboridae spiders have lost their venom apparatus while retaining toxin-like genes. The potent toxicity of their digestive fluids, a trait conserved across spiders, likely compensate for the absence of venom, ensuring effective prey immobilisation and digestion.

## Background

The arms race between predators and their prey is a fundamental driving force of animal adaptation. Over time, animals have evolved a range of strategies to succeed in this ongoing battle for survival and to gain advantages over their opponents. Among nature’s most effective weapons is venom - a potent cocktail of proteins and peptides capable of immobilising prey or deterring predators. Venomous animals are widespread across most branches of the animal tree of life [1,2]. This success can be attributed to the unique ability of venom to shift the battleground from the physical to the chemical level, enabling smaller and slower creatures to prevail over larger and faster ones. Moreover, venom has been linked to adaptive radiations in several animal clades, such as cone snails [3] and spiders [4–6], with diversification rates twice as high in venomous compared to non-venomous animal families [7].

The loss of venom might seem counterintuitive given its advantages, but it can be explained by its high metabolical cost due to the protein-rich nature of venom. Producing venom demands significant resources, creating competition with other physiological needs [6,8–10]. In species where venom use becomes less critical for survival, natural selection may favour its reduction or complete loss. Relaxed purifying selection can allow loss-of-function mutations to accumulate overtime through genetic drift, ultimately leading to the degradation of venom systems [11]. For example, in some species of catfish that rely on body size for predator defence, or sea snakes feeding exclusively on eggs [12], their venoms have become unnecessary [13]. In spiders, a similar phenomenon may have occurred in the Uloboridae family, which appears to lack functional venom glands [14]. In these spiders, the frontal part of their prosoma contains only muscle bundles where venom glands are typically located in araneomorph spiders. Whether uloborids are effectively venomless and how they subdue their prey without venom remains unclear.

Uloboridae spiders employ a unique hunting strategy characterised by extensive wrapping of prey in silk (up to hundreds of metres per prey), followed by the application of digestive fluids over the prey’s entire surface. This is in contrast to other spiders which typically regurgitated the fluids only on the area around their own mouthparts [15–18]. The broad application of the uloborid digestive fluids onto the prey appears to be lethal; Weng and colleagues [19] found that most ants in silk packages exposed to digestive fluids of the uloborid *Philoponella vicina* were dead, while most of the unwetted ants, or ants wetted with water, survived. This observation suggests that uloborids might have shifted the secretion of toxins from venom to digestive fluids, raising the question of whether they have retained venom components in other body secretions.

In this study, we tested the hypothesis that uloborids have replaced venom with toxic molecules in their digestive fluids, rendering venom glands redundant and leading to their evolutionary loss. We confirmed the absence of specialised venom-secreting glands in the prosoma of *Uloborus plumipes* and detected the expression of venom toxin homologs in various body tissues, particularly the midgut gland. Functional assays of midgut gland extracts showed no neurotoxicity against voltage-gated sodium or potassium channels. Nevertheless, they rapidly killed *Drosophila* flies, demonstrating the innately potency of uloborid digestive fluids. Similar results were observed for digestive fluids of the venomous Theridiidae common house spider, *Parasteatoda tepidariorum*, suggesting that the innovation in Uloboridae spiders lies not in the repurposing of toxins but rather in the strategic use of pre-existing secretions.

## Results

### Morphological evidence for the lack of venom glands

The objective of this study was to examine the structure of the chelicerae of *U. plumipes* and compare them with those of *P. tepidariorum*. Based on visual examination of dissected individuals and histological transverse sections across the chelicerae and anterior part of the prosoma, we confirmed the absence of venom glands in *U. plumipes* (Fig. 1). The chelicerae have a prominent epithelium and large muscles (Fig. 1A-E) but lack any kind of duct of the kind observed in *P. tepidariorum* (Fig. 1F, G).

**Figure 1:**
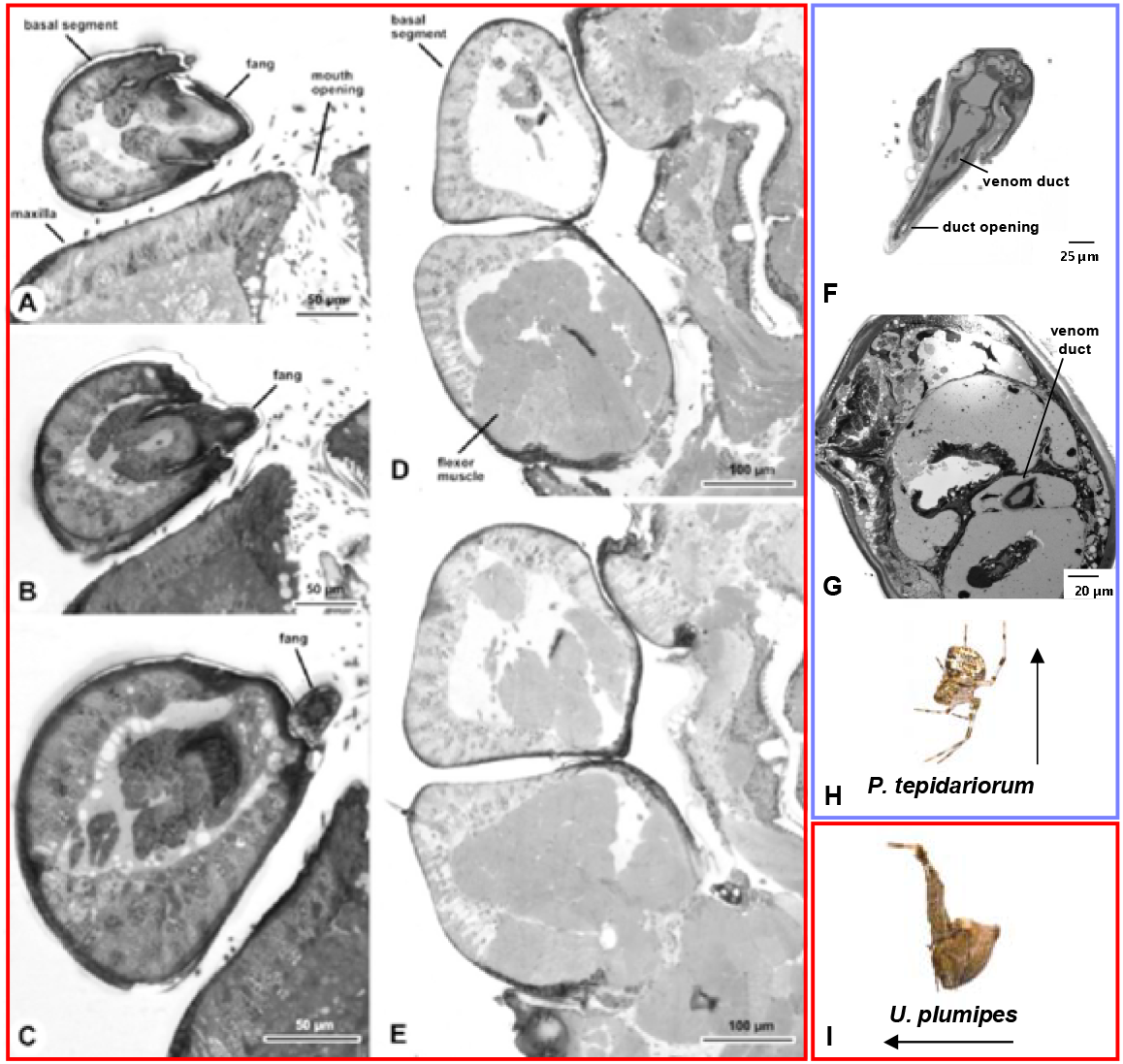
Histological transversal sections of the chelicerae of Uloborus plumipes and, for comparison, the venomous Parasteatoda tepidariorum. A-E: Cross-sections of *U. plumipes* (red), F-G: Cross-sections from *P. tepidariorum* (purple). A, B: Distal end of the chelicera and fang. C: Section through the area just above the fang base. Note that the fang does not contain any kind of canal. D, E: The proximal end of the chelicera, close to the transition to the prosoma, is filled with very large flexor muscles, but no duct nor venom gland base is visible. F: Venom duct and opening visible in the fang. G: Venom duct clearly visible in the chelicera. H: Image of *U. plumipes* by Olei under CC BY-SA 2.5 and I: Image of *P. tepidariorum* by J. Gallagher under CC-BY-2.0 via Wikimedia Commons.

Electron microscopy revealed the presence of three small pores at the tip of the uloborid fangs (Fig. 2A); however, compared to the much larger pore size of the theridiid fang (Fig. 2B), it is unlikely that the uloborid pores represent the opening of a venom duct. Similar small cuticular pits have been observed at the fang’s tip of the orb-web spider *Nephila clavata*, although their function remains unknown [20].

**Figure 2:**
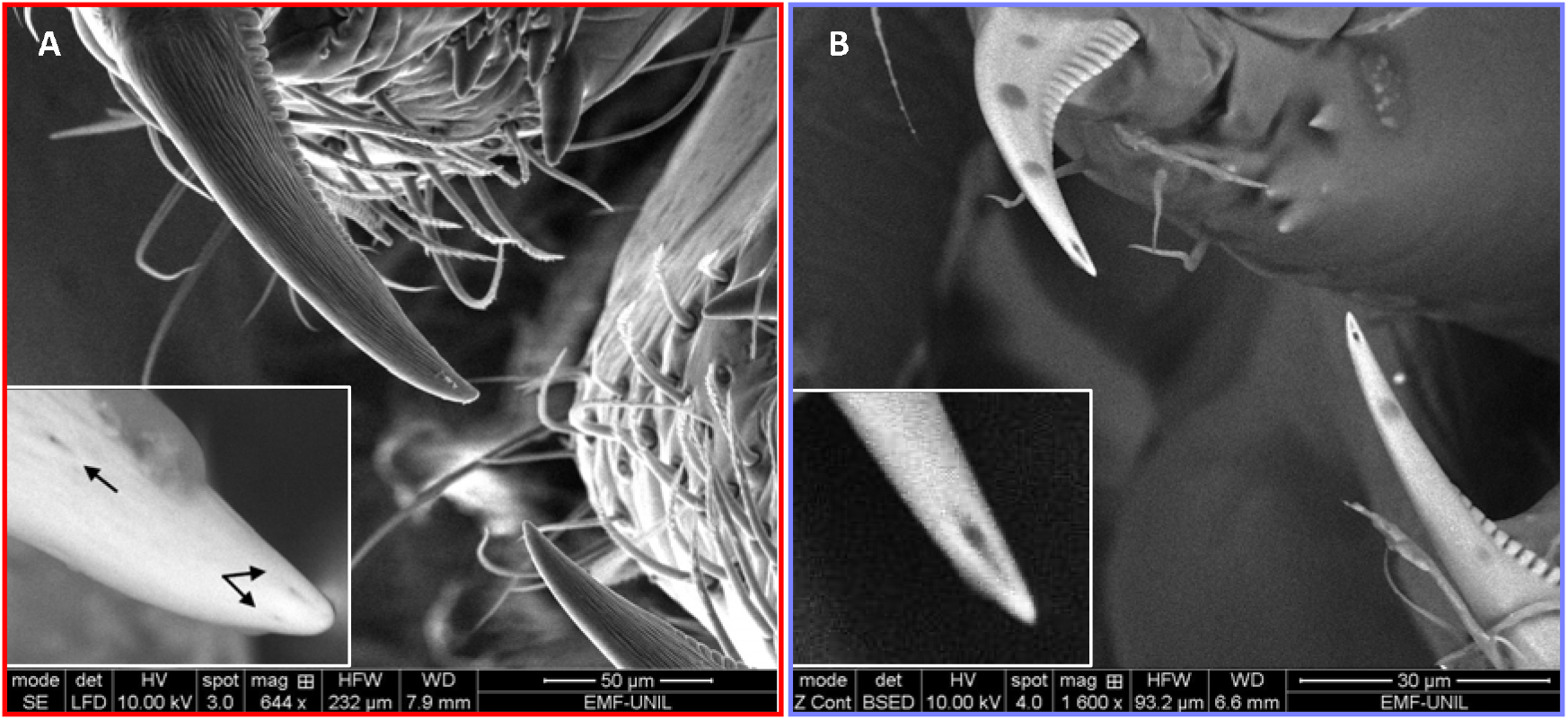
Fangs of Uloborus plumipes and Parasteatoda tepidariorum. A: *U. plumipes* possesses small pits at the tip of the fangs (black arrows in the zoomed insert). B: As a comparison, *P. tepidariorum* possesses a needle-like opening of the venom duct at the tip of the fangs.

### RNA-seq reveals the expression of putative toxins in the midgut gland of *U. plumipes*

#### De novo transcriptome assembly statistics

Although our morphological analysis confirmed the absence of a specialised venom-producing and delivery system in the anterior part of the spider’s body, it is possible that uloborids relocated the production of venom components to other organs that produce secretions that come into contact with the prey. To investigate this further, we performed RNA sequencing of 13 libraries from different tissues of *U. plumipes* including midgut gland, silk glands, chelicerae, prosoma, and gonads. The libraries ranged in size from 53 to 107 million reads (Supplementary Table S1).

The resulting *de novo* transcriptome contained 302’504 assembled transcripts, of which 20’243 (6.7%) passed our annotation pipeline (Table S2, Dataset 1). Our final assembly was 83.5% complete according to OMArk [21] (Fig. S1). OrthoVenn3 [22] identified 18’258 transcripts (90% of the total) as homologous to the Uloborus diversus genome annotation, and these were then clustered into 10’518 orthogroups. Among these orthogroups, 9’817 were shared with *U. diversus*.

The distribution of transcript counts was similar among libraries, except the midgut sample “UM1”, which had a smaller median value than the other libraries (Fig S2, S3). Principal component analysis showed that samples clustered by tissue type, with the first two components explaining 60% of the total variance (Fig S4).

#### Identification of toxin-like transcripts in the midgut gland

Manual screening of the final assembly identified 124 translated sequences matching venom-related proteins in the UniProt-ToxProt database [23] with biased expression across tissues (Dataset 2), and with 27 predicted to have neurotoxic activity using the NT-estimation model. The most represented gene families included astacin-like metalloprotease with 32 sequences (26%), serine proteases (13%), and cysteine-rich peptides such as atracotoxins, prokineticins, kunitz-type serine protease inhibitors, and thyroglobulin type-1 (Fig. 3A). Notably, most toxin-like transcripts were significantly upregulated in the midgut gland, supporting the hypothesis of venom components repurposed into digestive secretion (Fig. 3B).

**Figure 3:**
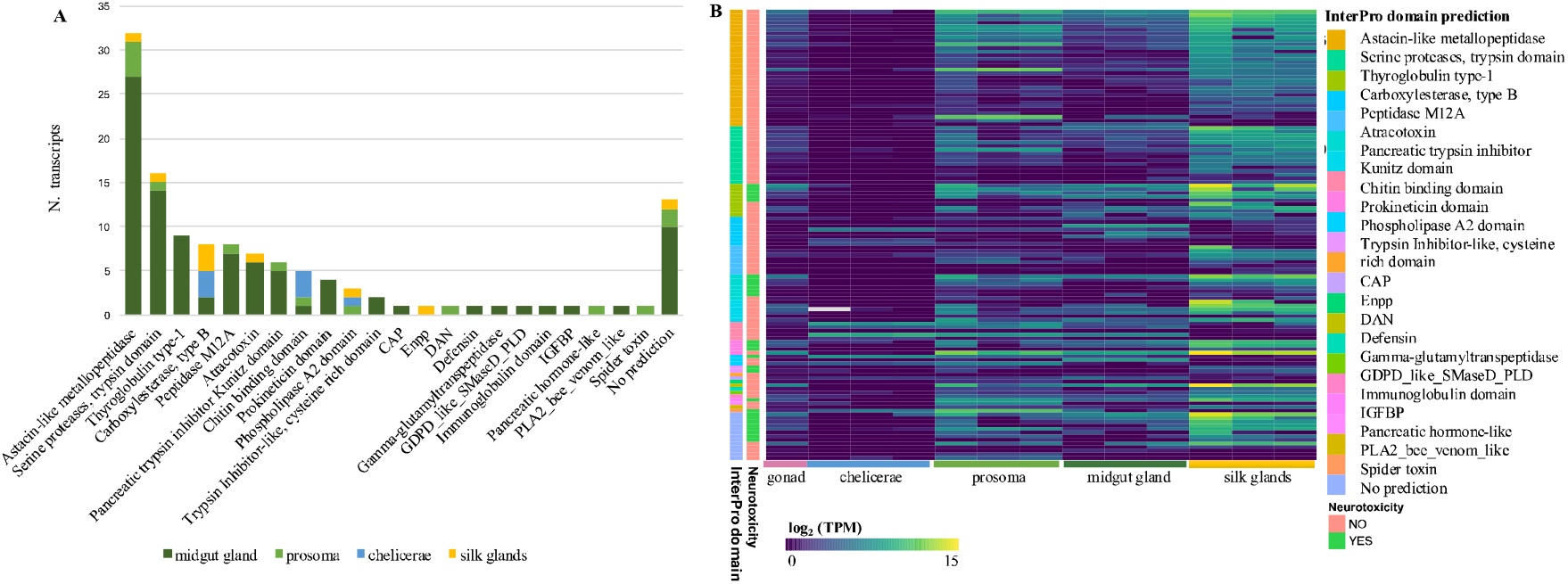
Classification and tissue distribution of *U. plumipes* toxin-like transcripts. A: Distribution of toxin-like transcripts classified based on their InterProScan predicted domains. The colour corresponds to the body tissue where their expression was significantly upregulated. ‘No prediction’ includes sequences without InterProScan annotation but with a positive BlastP hit. B) Expression levels of the 124 toxin-like transcripts across the tissues investigated. The transcripts are sorted by their InterPro domain as in (A), and neurotoxicity prediction is also reported.

Endopeptidases, such as metallo and serine proteases, constituted 43% of the upregulated transcripts in the midgut gland, consistent with their role in proteolysis and findings from prior proteomic studies [24–26]. While diverse, these transcripts were not the most highly expressed (Fig. 3b). Instead, the topmost expressed transcript was a cysteine-rich peptide orthologous to XP_054706256.1 in *U. diversus* (86% identity) with a prokineticin domain and a BlastP hit to U3-aranetoxin-Ce1a. Other highly expressed cysteine-rich peptides were predicted to be neurotoxic, with also BlastP hits to U3-araneotoxin as well as U24-ctenitoxins, the latter previously identified at the proteomic level in digestive fluids of *Uloborus* [26] and other spider species [24,25].

A notable toxin homolog expressed in the midgut gland was a Phospholipase D-like protein known as Sphingomyelinase D, a potent dermonecrotic component of *Loxosceles* venoms. Valladão et al. [26] confirmed the enzyme’s presence in the midgut gland, but as a β-type variant, which lack the severe necrotic activity observed in the α-type. However, the insecticidal potential of this β variant remains untested [26].

#### Defensin expression: an enigmatic toxin-like component

The second most highly expressed toxin-like transcript of *U. plumipes* was a defensin with a BlastP hit to the scorpion venom peptide BmKDfsin3 from Mesobuthus martensii (Fig. 4). Interestingly, this defensin was not annotated in the *U. diversus* genome. BlastN searches revealed the ortholog defensin gene on chromosome 4, while searches against the recently available *U. plumipes* genome identified four hits on four different contigs, one of which was 100% identical to our transcript.

**Figure 4:**
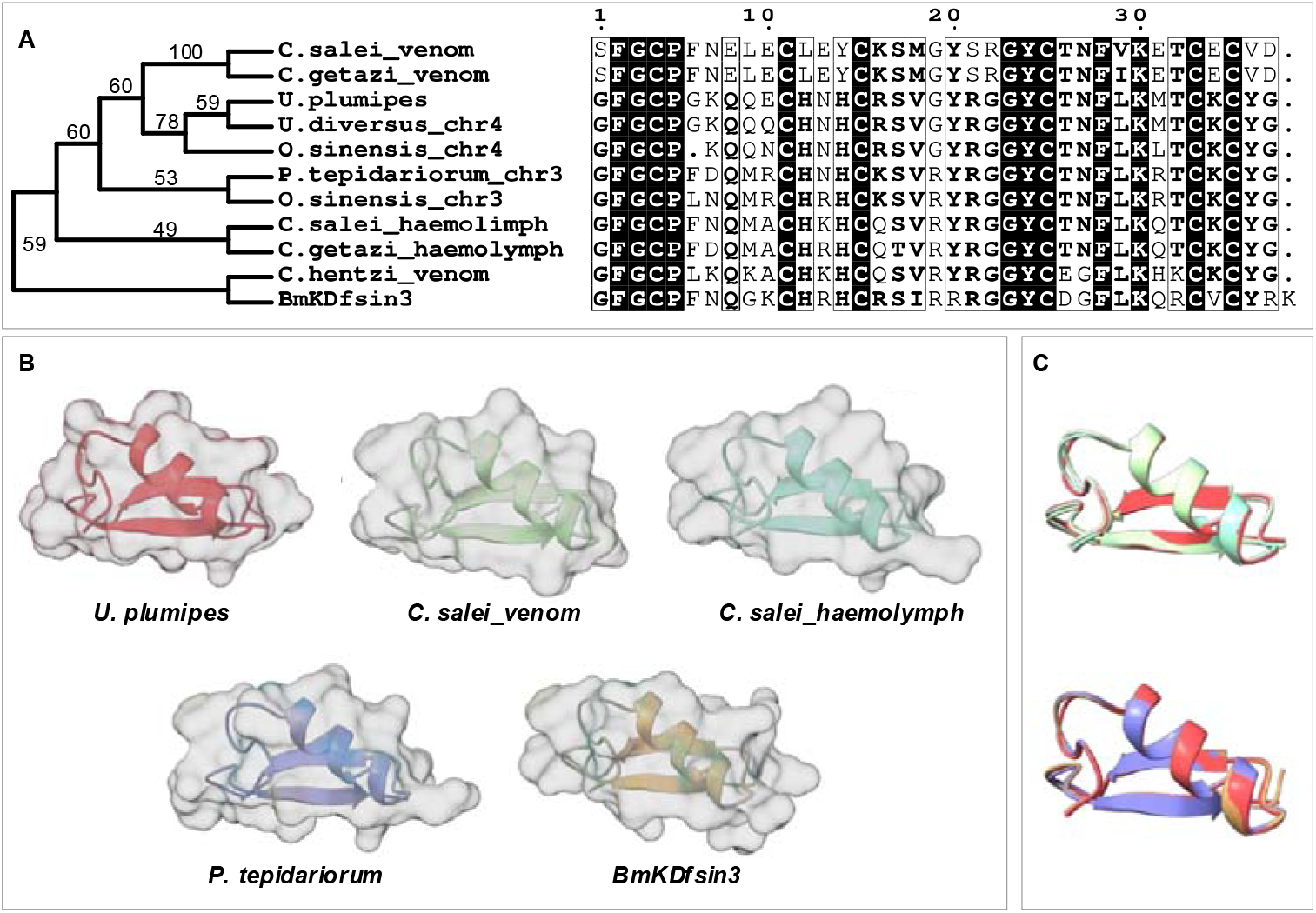
Phylogenetic analysis and 3D structure prediction of defensin. A: Neighbour-joining tree with branch bootstrap values and multiple sequence alignment of the mature region of the defensin found strongly upregulated in the midgut gland of *U. plumipes*. B: Alphafold 3 structure prediction of the defensins. Note the similarity between the copy in *U. plumipes* and that found in the venom glands of *C. salei*. C: Overlaid 3D structure predictions of defensins from *U. plumipes* and that from other species. The colour-code for the species is the same as in (B).

**Figure 5:**
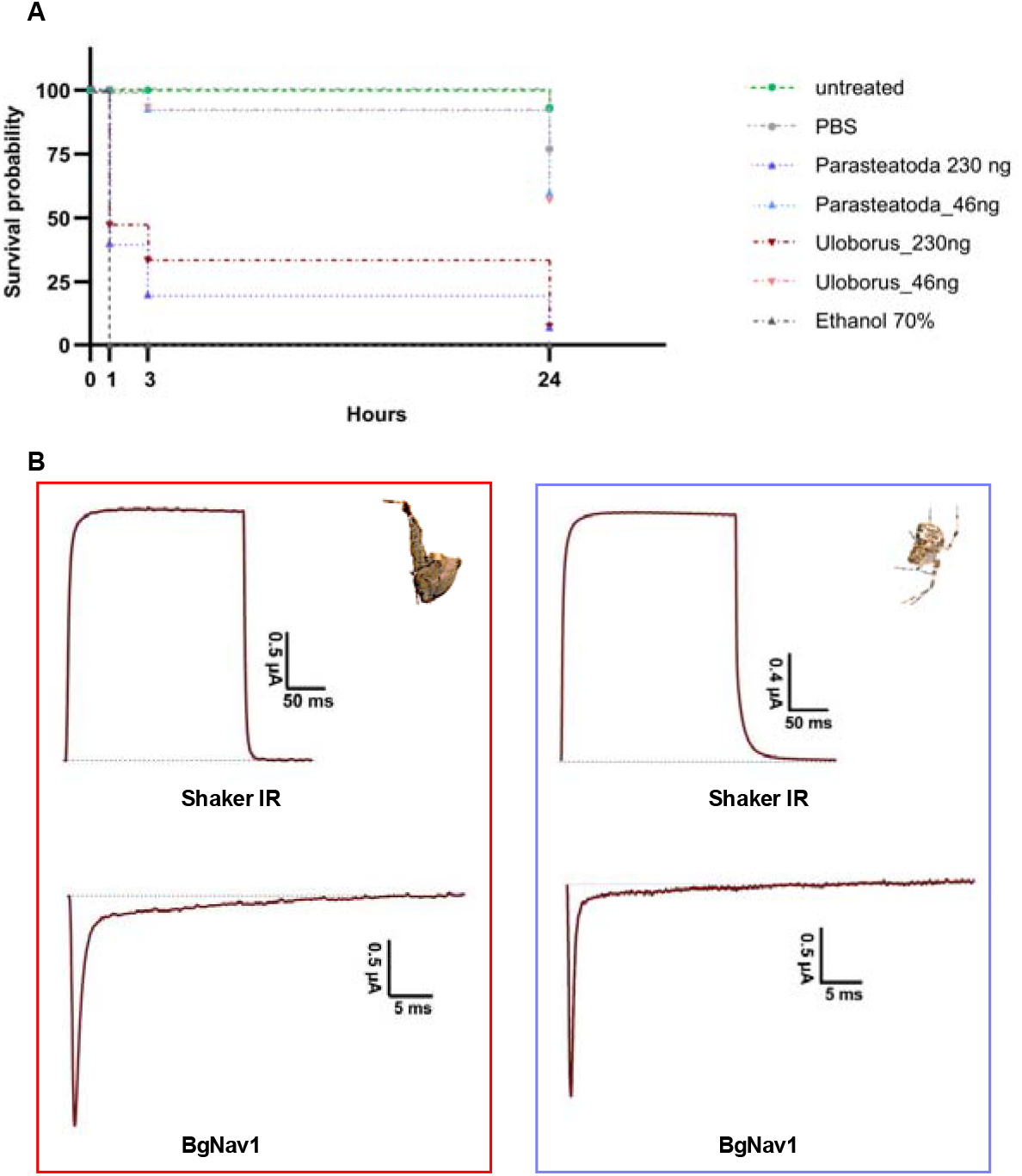
Bioassays of spider midgut gland extract. A: Survival rates of D. melanogaster injected with midgut gland extracts from *U. plumipes* and *P. tepidariorum* using 46nl of 5mg/ml (230ng) and 1mg/ml (46ng) of protein extract. The negative control is represented by the untreated group and flies injected with PBS. The positive control is represented by injections of 70% ethanol. B: Electrophysiological experiments with *U. plumipes* (red) and *P. tepidariorum* (purple) midgut gland extracts as representative traces of currents through the potassium (Shaker IR) and sodium (BgNav1) ion channels after application of 10 µg of extract.

Similarly, in another Uloboridae species, *Octonoba sinensis* [27], three defensin genes were found: the ortholog of *U. plumipes* on chromosome 4, a second gene on the same chromosome with less identity, and another on chromosome 3. The ortholog on chromosome 4 is embedded within the gene “g1332” which seems to have been misannotated since its coding sequence does not match any spider genes except for a retrotransposon in *Araneus ventricosus* with 32% identity. Mapping available SRA libraries from multiple *O. sinensis* tissues to the added defensin annotation showed high expression in the ‘abdomen’ sample (Fig. S5). It remains uncertain if the midgut gland was dissected within the abdomen sample, although the expression pattern suggests that it likely was. Beyond Uloboridae, a defensin gene is present on chromosome 3 in *P. tepidariorum*, as well as in other spider genomes. Notably, defensins were found in the venom glands of *Cupiennius* species, distinct from those expressed in the haemolymph [28]. Phylogenetic analysis and Alphafold 3 [29] predictions of the mature region of the defensin sequence suggest two copies (Fig. 4A): one likely expressed in haemolymph on chromosome 3 and another more derived on chromosome 4, highly expressed in the midgut gland of Uloboridae spiders, resembling the venom gland defensin of *Cupiennius* (Fig. 4B, C).

#### Toxin-like transcripts in other body parts

Outside the midgut gland, toxin-like transcripts were less highly expressed, although significantly upregulated to a specific body part. For instance, a U8-agatoxin-Ao1a-like sequence with a knottin domain, which is one of the most abundant domains in spider venoms [6], was highly expressed in the prosoma. However, it presented a disruption in the canonical cysteine framework (C-CXC-CC-CXC-CXC-C), with the absence of the second and tenth cysteine residues (Fig. S6).

In the silk glands, we detected various acetylcholinesterase genes. Acetylcholinesterase is an enzyme family typically involved in neurotransmitter breakdown at postsynaptic neuromuscular junctions. While its presence in the venoms of spiders, centipedes, and other organisms has been noted, its role in spider venom, as well as in the silk, remains uncertain. Nevertheless, it has been hypothesized to serve a trophic function [30].

Finally, several ferritin transcripts ranked among the most highly expressed in the whole transcriptome. Ferritin, commonly reported in cone snail venom glands [31,32], has also been detected in spider digestive fluids [24], suggesting a broader physiological role.

### Midgut gland extracts are insecticidal

The upregulation of toxin-like transcripts in the midgut gland, particularly of peptides with predicted neurotoxic activity, supports the hypothesis that Uloboridae spiders have shifted venom-like components into their digestive fluids to aid in prey immobilisation. Since Uloboridae do not inject venom, these potent digestive fluids may play a central role in incapacitating prey. To test this hypothesis, we compared the midgut gland extracts of *U. plumipes* and *P. tepidariorum* by injecting them into *Drosophila suzukii*.

Both extracts were highly insecticidal, with 230 ng killing over 50% of flies within one-hour post-injection. Even as little as 46 ng killed approximately 40% of flies within 24h, with *U. plumipes* extracts slightly more potent, though not statistically significant (t = -0.25, df = 3.9, p = 0.4) (Fig. 4A).

### Midgut gland extract does not block sodium and potassium channels

Considering the high expression of transcripts with predicted neurotoxic activity, we tested whether the midgut extract’s insecticidal effect was due to neurotoxins targeting sodium and potassium channels, which are among the most common voltage-gated ion channel targets of spider venoms [6]. Using a two-electrode voltage clamp assay, we examined whether 10 µg of extract inhibited sodium channels from *Blatella germanica* (BgNav1) or potassium channels from *D. melanogaster* (Shaker IR).

The extracts showed no inhibitory effects on either channel (Fig. 4B). Furthermore, even at concentrations up to 30 µg, the extracts did not induce cytolysis in oocytes, indicating that the midgut secretions do not cause membrane damage or cell integrity disruption, as pore-forming toxins do.

## Discussion

Spiders are efficient predators, relying on foraging webs and potent venoms to capture and subdue prey. Nearly all spiders inject venom, with spiders of the Uloboridae family being a remarkable exception. For nearly a century, uloborids were believed to lack venom glands, a notion originating from an early anatomical sketch [14]. However, this assumption had remained largely unexamined until now. In this work, we investigated whether uloborids have indeed lost their venom apparatus and whether venom-associated toxins were repurposed in other secretions.

### Uloboridae spiders have effectively lost their venom apparatus but not their toxins

Our histological analyses confirmed the absence of venom glands in the prosoma of *U. plumipes*. Furthermore, the venom duct opening at the fang tip was absent, indicating complete loss of the venom system. Interestingly, we observed small pores on the fangs, a feature also noted in other spiders [20], though their function remains unclear. While histology confirmed the absence of venom structures, our transcriptomic analysis revealed a more complex story: venom-like components are still be present and expressed, specifically in the midgut gland, the organ secreting digestive fluids. A proteomic study on a related species confirmed the presence of similar toxins [26], and our functional assays demonstrated the lethal effects of midgut gland extracts on *Drosophila* flies. Even at low concentrations, these extracts were highly insecticidal, underscoring their potency. This discovery suggests that uloborids may have repurposed venom toxins, integrating them into their digestive system for prey immobilisation.

### Toxins in spider digestive fluids: an evolutionary adaptation?

The presence of venom components in digestive fluids is not unique to Uloboridae. Studies on venomous species of different spider taxa, such as the golden orb-weaver Nephila cruentata [24], or the Brazilian white-knee tarantula *Acanthoscurria geniculata* and the velvet spider *Stegodyphus mimosarum* [25], have detected similar proteins in their digestive secretions, albeit in varying proportions. For instance, 66% of venom components in *S. mimosarum* were also found in digestive fluids, compared to only 9% in *A. geniculata* [25]. Conversely, digestive enzymes have been detected in venoms, possibly starting the extra-oral digestion [25]. Why toxins exist in digestive fluids remains unclear, with no definitive hypotheses proposed thus far.

We suggest that this dual presence serves as an evolutionary safety mechanism, ensuring prey immobilisation when venom is inadequate. In large spiders such as *A. geniculata*, which has strong jaws and can kill prey just with the strength of their bite alone, fewer toxins are found in the digestive fluids. While their ‘safety mechanism’ is brute force, smaller spiders may rely on a dual chemical weapon. Our functional assay demonstrated that also the midgut extract of the venomous *P. tepidariorum* was equally potent to that of *U. plumipes* in killing *Drosophila* flies.

### How uloborids kill their prey

We propose that Uloboridae spiders have not evolved novel toxic secretions but rather altered how existing secretions are deployed. Unlike typical spiders, which apply digestive fluids after venom injection and only near their mouthparts, uloborids exhibit a unique feeding behaviour. They wrap their prey extensively in silk and cover the entire body with digestive fluids [15–18]. It is only after the application of digestive fluids that prey death occurs [19]. We can think of this external application of toxin secretion as mimicking the internal diffusion of venom. Proteolytic enzymes, such as astacin-like metalloproteases and serine proteases, degrade intersegmental membranes, acting as spreading factors and facilitating toxin entry or potentially causing death through enzymatic activity alone [33].

A key question remains: do uloborids digestive secretion contain neurotoxins capable of paralysing prey? And if so, are these toxins unique to Uloboridae, or shared with venomous species?

While our transcriptomic analysis of *U. plumipes* revealed upregulation of genes with predicted neurotoxic activity in the midgut gland, their exact role remains speculative. Functional assays on sodium (Shaker IR) and potassium (BgNav1) channels showed no inhibition of these commonly targeted voltage-gated ion channels. However, many potential targets, such as calcium channels, acid-sensing ion channels, or glutamate receptors, remain untested [6]. Furthermore, toxin efficacy often depends on isoform specificity. For example, scorpion toxin MeuTXKα1 effectively blocks rKv1.1, hKv1.3, and *Shaker* IR channels but not other isoforms such as rKv1.2, rKv1.4, rKv1.5, rKv1.6, or hERG [34]. This suggests that uloborid neurotoxins may act on yet unidentified targets, necessitating further exploration.

### A promising candidate: defensin

Among the identified candidate toxins, defensin stood out due to its high expression, particularly in the midgut gland. Defensins are small peptides with antimicrobial properties found in various organisms, including plants, arthropods, and vertebrates. In scorpions, defensins have dual roles: they exhibit antimicrobial activity and block potassium channels (Kv1.1, Kv1.2, Kv1.3 and SK3), functioning similarly to neurotoxins like OSK1 and ScyTx [35]. Notably, slight modifications can transform an insect defensin into a potent scorpion-like potassium channel inhibitor [36]. Despite the translated sequence of *U. plumipes* defensin being predicted neurotoxic, our electrophysiology experiment showed no inhibition on the BgNav1 channel. However, it is possible that other ion channels are targeted, as discussed earlier.

Beyond scorpions, defensins have also been detected in the venom glands of the bromeliad spider species *C. salei* and *C. getazi* [28]. Our genomic and phylogenetic analyses show that, after duplication, the copy on chromosome 4 underwent upregulation in the midgut gland of Uloboridae and the venom glands of *Cupiennius*, while the copy on chromosome 3 appears to maintain the original immunity role. Further genomic studies using the newly available spider genomes could provide more clarity on the evolutionary history of this intriguing gene family. Our BlastN searches also highlight the challenges in annotating these small peptides, which may hinder proteomic analyses. Defensins have not been reported in the digestive fluids of various spiders [25], including *Uloborus* sp. [26], and this absence may reflect technical limitations due to the lack of the defensin sequence in the databases used for peptide identification rather than a true absence in secretion.

## Conclusions

This study investigated the evolutionary loss of venom in the spider *U. plumipes*, confirming the absence of a venom apparatus in Uloboridae while revealing the high expression of venom toxins in the midgut gland. These findings demonstrate that spider toxins are not exclusively confined to specialised venom-secreting glands but are also play a role in the digestive system. This supports an evolutionary link between the two systems, suggesting that toxins may have initially served digestive functions before being co-opted for venom use. In venomous spiders, toxins in the digestive fluids likely act as a secondary mechanism for effective prey immobilisation, while Uloboridae have adapted this trait as their primary strategy by dispersing toxic fluids across the prey’s body. This study highlights the complexity of spider biology and emphasises the importance of investigating toxin expression across tissues to fully understand their evolutionary and functional roles.

## Methods

### Specimens

Individuals of *Uloborus plumipes* were collected from greenhouses in Viernheim and Merzig in Germany, and in Lausanne, Switzerland. *Parasteatoda tepidariorum* spiders are from a laboratory colony which is kept at 27°C with 70% humidity on a gradual light/dark cycle of 16/8 hours and fed twice a week with Musca domestica flies.

### Morphological and histological analyses

To verify the absence of venom-producing glands in *U. plumipes* we performed a histological analysis of the chelicerae and anterior part of the prosoma. We dissected five individuals of *U. plumipes* directly in the fixative, Karnovsky’s solution [37], and fixed the chelicerae with part of the prosoma overnight. After washing in 0.1M phosphate buffer, the samples were post-fixed in 2% osmium tetroxide solution for two hours and dehydrated using a graded series of ethanol. Embedding was carried out using Embed812 resin embedding kit (Science Services GmbH, München, Germany). During the final step, samples were transferred into a “VacuTherm” vacuum heating cabinet (Thermo Fisher Scientific, Waltham, Massachusetts, USA) and incubated at 40 °C and 100 mbar for 3 × 30 min. Polymerization of the resin blocks was carried out in a heating cabinet at 60 °C for a minimum of 24 h. Semi-thin sections were obtained with a Leica UC6 ultra-microtome (Leica Microsystems GmbH, Wetzlar, Germany), with a DiATOME histo Jumbo diamond knife (Diatome Ltd., Nidau, Switzerland) at a thickness of 700 nm. Staining was done with toluidine blue at 70 °C, and images were obtained with a Fritz Slide Scanner (PreciPoint GmbH, Freising, Germany).

Individuals of *P. tepidariorum* were fixed in 2.5% glutaraldehyde for 1 hour at room temperature and subsequently fixed overnight at 4°C. After several washes with 0.1 M phosphate buffer, the samples were postfixed in 2% osmium tetroxide solution for 1 hour at room temperature, then washed with water, dehydrated through a graded series of acetone and subsequently a graded series of resin, and finally embedded using Spurr Low-Viscosity Embedding kit (SIGMA). The embedded spiders were incubated in resin blocks at 60°C for 48 hours. Semi-thin sections were obtained with a Leica EM UC7 Ultramicrotome (Leica Microsystem), with a DiATOME diamond knife (Diatome Ltd., Switzerland) at 700 nm thickness. Sections were stained with 0.5% toluidine blue, and images were acquired using Zeiss Axio Imager Z2 (Leica Microsystem).

### Scanning electron microscopy

We used three individuals to examine the fang morphology. Specifically, we aimed at verifying whether the opening of the venom duct at the tip of the fangs were present, or whether they disappeared as a consequence of the loss of the venom-secreting apparatus. The spiders were fixed in liquid nitrogen, placed on an aluminium holder with double-sided carbon adhesive, and examined with a scanning electron microscope (Quanta FEG 250, TFS) using the environmental mode (partial pressure 80Pa). The detectors used were the Large Field Detector and the Backscattered electron detector (BSED) at 10kV spot 4, working distance between 7.9 and 6.6mm.

### RNA-seq

Tissue samples of chelicerae, prosoma, midgut gland, silk gland and gonads (ovaries) were dissected from approximately 12 adult individuals. To obtain enough RNA, for each tissue, we pooled multiple samples for a total of three replicates except the ovaries for which we had only one. Total RNA was isolated using the TRIzol™ Plus RNA Purification kit (ThermoFisher) following manufacturer’s instructions, with an additional on-column DNA purification step. Thirteen cDNA libraries were generated with the TruSeq RNA Sample Preparation kit (Illumina) with 150 read length, followed by pair-end sequencing on an Illumina NovaSeq at the Genomic Technologies Facility of the University of Lausanne, Switzerland.

### Assembly and annotation

Raw reads were assessed with FastQC v0.11.9 [38] and quality-filtered with Fastp v0.22.0 [39]. Reads shorter than 30bp were discarded. As no genome for Uloboridae spiders was available at time of the project, all reads from all tissues were concatenated and used for de novo assembly with SPAdes v3.15.3 [40]. We adopted a multi-step quality filtering approach to prune the raw transcriptome assembly. First, we used the program Borf v1.2 [41] to predict open reading frames (ORFs) and retained only sequences with a minimum length of 20 amino acids and a complete ORF. Next, we annotated the filtered amino acid sequences with BlastP [42] against multiple databases, including the NCBI non-redundant pre-formatted Refseq database, UniProt/SwissProt [43], UniProt-ToxProt [23], Arachnoserver [44], and a customised database consisting of 11 spider genomes (Supplementary Table S3). All databases were downloaded on July 4^th^, 2022. Protein families and domains were predicted with InterProScan v5.51.85.0 [45], and signal peptides were detected using SignalP-6.0 [46]. Only transcripts with a BlastP hit to at least one database and e-value < 1e-5 were retained. We further reduced redundancy by clustering all nucleotide sequences with > 99% identity using CD-HIT v4.8.1 (Fu et al. 2012). Orthologous clusters with *Uloborus diversus* (GCF_026930045) genes [47] were identified using the web platform OrthoVenn3 [22]. The completeness of the final assembled transcriptome was assessed by OMArk [21] by calculating the overlap of the non-redundant annotated genes and conserved ancestral gene set of the phylum Arthropod.

### Toxin-like transcripts identification

We assigned a transcript as toxin-like if its best hit, defined as having the highest bitscore, was a protein listed in the UniProt-ToxProt database [23]. Furthermore, based on the assumption that toxins should have biased expression specifically to the gland that secretes them, we further minimise the chances of false positives by running a differential expression analysis and keeping only those toxin-like transcripts which were significantly upregulated (see below). Presence of a knottin domain was tested with the tool Knotter 1D from the Knottin database website [48], and neurotoxic activity was predicted using NT_estimation, a deep learning approach which uses a peptide data augmentation method to improves the recognition of spider neurotoxic peptides via a convolutional neural network model (Lee et al. 2021). The putative toxins were classified into their corresponding protein families based on their predicted InterPro protein domain [45]

### Expression level quantification

Transcript abundances were quantified using Kallisto v0.48.0 (Bray et al., 2016) with default parameters for paired-end reads. Downstream analyses were conducted in R version 4.1.3 [49]. Count distribution across libraries was inspected with the package *vioplot* v 0.3.7 (Adler et al., 2021) and genes with TPM value >= 1 in at least one library were kept for further analyses. Consistency of expression patterns between samples from the same tissue was assessed by means of principal component analysis, and the percentage of explained variance by each component was calculated using the function fviz_eig in Ade4 v1.7.19 (Dray and Dufour, 2007).

Differential expression analysis was performed in Sleuth v0.30.0 (Pimentel et al., 2017) and the fold change (FC) of transcripts was calculated as the ratio between the highest and the second highest expression values. Only putative toxin transcripts with fold change >= 2 and q-val < 0.05 were retained.

Because *Sleuth* automatically filter out genes which are exclusively expressed in one tissue type, we also checked those which did not pass Sleuth filter and kept all the sequences with FC >= 2.

### Genomic investigation of defensin

Considering the high expression levels of defensin in the midgut gland, and the lack of ortholog with *U. diversus*, we thought to investigate further this putative toxin in other spider species, specifically other Uloboridae whose genome became available during the course of the study. First, we performed BlastN [42] searches against the whole genome of the Ulobordae *U. plumipes, U. diversus*, and *Octonoba sinensis*, as well as *P. tepidariorum* and the NCBI non-redundant database. We used SeaView v5.0.4 [50] for multisequence alignment with MUSCLE v3.8.31 [51] and to obtain a Neighbour-Joining tree with bootstrap from 100 replicates. Three-dimensional (3D) structures were predicted using Alphafold v3 [29]. Expression levels were examined for those species with multi-tissue RNA-Seq data, i.e. *O. sinensis* and *P. tepidariorum*. For the latter, we used the expression quantification from Hassan et al. [52], while for *O. sinensis* we aligned reads from a subset of SRA (see Table S4) using STAR v2.7.10b [53] and quantified transcript abundances with Kallisto v0.48.0 [54] after adding the defensin transcripts to the annotation. As the defensin was embedded into gene “g1332” which seem to be spurious annotation, we performed the quantification also after removing this gene.

### Midgut gland protein extraction and *in vivo* experiments

Dissected midgut gland tissues from *U. plumipes* and *P. tepidariorum* were physically disrupted with a Potter-Elvehjem homogeniser as previously described by Valladão et al. [26] Briefly, samples were homogenized in a falcon tube containing 500µL of water and then centrifuged at 16.000g for 30 min at 4°C. The supernatant containing the digestive fluids was collected and lyophilised using a Beta 2-8 LSCplus (Christ) freeze drier. Lyophilised midgut extracts were used for downstream experiments. Protein content was quantified using a BCA-Protein-Assay kit (Merck).

Insecticidal activity was assessed *in vivo* as described earlier [55]. For the injections, concentrations of 5 mg/mL and 1 mg/mL of the extracts were prepared in PBS. *Drosophila suzukii* flies from a laboratory stock, originally sourced from Ontario, Canada, were reared in a climate chamber at 26 °C with 60% humidity and a 12 h photoperiod. They were fed on 10.8% soybean and cornmeal mix, 0.8% agar, 8% malt, 2.2% molasses, 1% nipagin and 0.625% propionic acid media. For the insecticidal assay, we used 4–6-day old adult flies. Flies were anaesthetized on a CO_2_-Pad (Inject+Matic). Volumes of 46 nl were injected intrathoracically using glass capillaries pulled on a P-2000 Laser-Based Micropipette Puller (Sutter Instrument) held on a Nanoject II device (Drummond Scientific). Injections were performed under a Stemi 508 Stereomicroscope (Zeiss). Negative controls consisted of untreaded flies and flies injected with 46 nl of PBS, while positive controls were injected with 46 nl of 70% ethanol. Batches of 10 flies then incubated in ∅ 29 × 95 mm vials with foam stoppers (Nerbe Plus) filled 1/16 with food media. Survival rates and signs of paralysis were assessed 1h, 3h and 24h post injection.

### Two-electrode voltage clamp electrophysiology

We investigated neurotoxicity of the digestive fluids by examining whether 10 µg of midgut extract would inhibit potassium and the sodium channels. Recordings were performed at room temperature (18–22 °C) using a Geneclamp 500 amplifier (Molecular Devices, San Jose, CA, USA) controlled by a pClamp data acquisition system (Axon Instruments, San Jose, CA USA). Whole-cell currents from *Xenopus laevis* oocytes were recorded 1–4 days after cRNA injection. The bath solution composition was ND96. Voltage and current electrodes were filled with 3 M KCl. Resistances of both electrodes were kept at 0.7–1.5 MΩ. Elicited currents were sampled at 1 kHz and filtered at 0.5 kHz (for potassium currents) or sampled at 20 kHz and filtered at 2 kHz (for sodium currents) using a four-pole low-pass Bessel filter. Leak subtraction was performed using a −P/4 protocol.

For the electrophysiological measurements, we used a 2 s ramp protocol from −120 to +80 mV, with cells clamped at −120 mV holding potential. Shaker IR currents were evoked by 500 ms step depolarizations to 0 mV, followed by a 500 ms pulse to −50 mV from a holding potential of −90 mV. Sodium current traces were evoked by a 100 ms depolarization to 0 mV. All data were obtained in at least five independent experiments (n ≥ 5). Non-injected oocytes without expressing any type of ion channel were used to check whether the midgut extracts were cytotoxic to oocytes. Animal experiments using X. laevis were approved by the Animal Ethics Committee of KU Leuven in accordance with EU Council Directive 2010/63/EU.

### Expression of ion channels in *X. Laevis* oocytes

The following genes encoding ion channel subunits were expressed in X. laevis oocytes: the voltage-gated potassium channel, Shaker-IR from *Drosophila* melanogaster and the voltage-gated sodium channel BgNav1 from *Blattella germanica*. Linearised plasmids bearing the ion channel genes were transcribed using the mMESSAGE mMACHINE T7 transcription kits (Ambion, Austin, TX, USA) to prepare the respective cRNA. The harvesting of stage V–VI oocytes from anesthetized female *X. laevis* frogs was described previously [56]. Oocytes were injected with 50 nl of cRNA at a concentration of 1 ng/nl using a micro-injector (Drummond Scientific, Broomall, PA, USA). The oocytes were incubated at 16°C in ND96 solution containing (in mM): NaCl, 96; KCl, 2; CaCl2, 1.8; MgCl2, 2; and HEPES, 5 (pH 7.4), supplemented with 50 mg/l gentamicin sulfate.

## Acknowledgements

This project has received funding from the European Union’s Horizon 2020 research and innovation programme under Marie Skłodowska-Curie Grant Agreement 845674 to G.Z. and the Etat de Vaud to M.R-R. We are grateful to Daniel Gritz and Vanessa Oehmig from the German Arachnological Society (DeArGe) who provided specimens of *U. plumipes* to our study. We thank J. Marquis from GTF for support with library preparation and sequencing.

## Author contributions

X.P. performed the RNA extraction and the transcriptome annotation and analysis. L.D., J.D, and T.L. performed the *in vivo* experiments. T.D., P.M., and A.H. performed the morphological and histological analyses. S.P. and J.T. performed the electrophysiology experiments. A.M. performed the electron microscopy. M.R-R. contributed to the transcriptomic analysis. G.Z. and T.L. conceived the study. G.Z. contributed to the transcriptomic analysis and wrote the manuscript, with input from all authors.

## References

1. Schendel V, Rash LD, Jenner RA, Undheims EAB. 2019 The diversity of venom: The importance of behavior and venom system morphology in understanding its ecology and evolution. Toxins 11, 666. (doi:10.3390/toxins11110666)

2. Modica MV et al. 2021 The new COST Action European Venom Network (EUVEN)— synergy and future perspectives of modern venomics. GigaScience 10, giab019. (doi:10.1093/gigascience/giab019)

3. Puillandre N, Fedosov AE, Kantor YI. 2017 Systematics and evolution of the Conoidea. In Evolution of Venomous Animals and Their Toxins (ed A Malhotra), pp. 367–398. Dordrecht: Springer Netherlands. (doi:10.1007/978-94-007-6458-3_19)

4. Vassilevski AA, Kozlov SA, Grishin EV. 2009 Molecular diversity of spider venom. Biochem. Mosc. 74, 1505–1534. (doi:10.1134/S0006297909130069)

5. King GF, Hardy MC. 2013 Spider-venom peptides: Structure, pharmacology, and potential for control of insect pests. Annu. Rev. Entomol. 58, 475–496. (doi:10.1146/annurev-ento-120811-153650)

6. Lüddecke T, Herzig V, von Reumont BM, Vilcinskas A. 2021 The biology and evolution of spider venoms. Biol. Rev. 97, 163–178. (doi:10.1111/brv.12793)

7. Arbuckle K, Harris RJ. 2021 Radiating pain: Venom has contributed to the diversification of the largest radiations of vertebrate and invertebrate animals. BMC Ecol. Evol. 21, 150. (doi:10.1186/s12862-021-01880-z)

8. Nisani Z, Hayes WK. 2011 Defensive stinging by Parabuthus transvaalicus scorpions: Risk assessment and venom metering. Anim. Behav. 81, 627–633. (doi:10.1016/j.anbehav.2010.12.010)

9. Nelsen DR, Kelln W, Hayes WK. 2014 Poke but don’t pinch: Risk assessment and venom metering in the western black widow spider, Latrodectus hesperus. Anim. Behav. 89, 107–114. (doi:10.1016/j.anbehav.2013.12.019)

10. Cooper AM, Nelsen DR, Hayes WK. 2017 The Strategic use of venom by spiders. In Evolution Of Venomous Animals And Their Toxins (ed A Malhotra), pp. 145–166. Dordrecht: Springer Netherlands. (doi:10.1007/978-94-007-6458-3_13)

11. Morgenstern D, King GF. 2013 The venom optimization hypothesis revisited. Toxicon 63, 120–128. (doi:10.1016/j.toxicon.2012.11.022)

12. Li M, Fry BG, Kini RM. 2005 Eggs-only diet: Its implications for the toxin profile changes and ecology of the marbled sea snake (Aipysurus eydouxii). J. Mol. Evol. 60, 81–89. (doi:10.1007/s00239-004-0138-0)

13. Wright JJ. 2017 Evolutionary history of venom glands in the Siluriformes. In Evolution of Venomous Animals and Their Toxins (ed A Malhotra), pp. 279–301. Dordrecht: Springer Netherlands. (doi:10.1007/978-94-007-6458-3_9)

14. Millot J. 1931 Les glandes venimeuses des Araneides. Ann. Sci. Nat. Zool. 14, 113–147.

15. Lubin YD. 1986 Web building and prey capture in the Uloboridae. SpidersWebs Behav. Evol., 132–171.

16. Opell BD. 1988 Prey handling and food extraction by the triangle-web spider Hyptiotes cavatus (Uloboridae). J. Arachnol. 16, 272–274.

17. Eberhard WG, Barrantes G, Weng J-L. 2006 The mystery of how spiders extract food without masticating prey. Bull.-Br. Arachnol. Soc. 13, 372.

18. Eberhard WG, Barrantes G, Weng J-L. 2006 Tie them up tight: wrapping by Philoponella vicina spiders breaks, compresses and sometimes kills their prey. Naturwissenschaften 93, x251–254. (doi:10.1007/s00114-006-0094-1)

19. Weng J-L, Barrantes G, Eberhard WG. 2006 Feeding by Philoponella vicina (Araneae, Uloboridae) and how uloborid spiders lost their venom glands. Can. J. Zool. 84, 1752– 1762. (doi:10.1139/z06-149)

20. Moon M-J, Yu M-H. 2007 Fine structure of the chelicera in the spider Nephila clavata. Entomol. Res. 37, 167–172. (doi:10.1111/j.1748-5967.2007.00108.x)

21. Nevers Y, Rossier V, Train CM, Altenhoff A, Dessimoz C, Glover N. 2022 Multifaceted quality assessment of gene repertoire annotation with OMArk. Nat. Biotechnol., 2022.11.25.517970. (doi:10.1101/2022.11.25.517970)

22. Sun J, Lu F, Luo Y, Bie L, Xu L, Wang Y. 2023 OrthoVenn3: an integrated platform for exploring and visualizing orthologous data across genomes. Nucleic Acids Res. 51, W397–W403. (doi:10.1093/nar/gkad313)

23. Jungo F, Bougueleret L, Xenarios I, Poux S. 2012 The UniProtKB/Swiss-Prot Tox-Prot program: A central hub of integrated venom protein data. Toxicon 60, 551–557. (doi:10.1016/j.toxicon.2012.03.010)

24. Fuzita FJ, Pinkse MWH, Patane JSL, Verhaert PDEM, Lopes AR. 2016 High throughput techniques to reveal the molecular physiology and evolution of digestion in spiders. BMC Genomics 17, 716. (doi:10.1186/s12864-016-3048-9)

25. Walter A, Bechsgaard J, Scavenius C, Dyrlund TS, Sanggaard KW, Enghild JJ, Bilde T. 2017 Characterisation of protein families in spider digestive fluids and their role in extra-oral digestion. BMC Genomics 18, 600. (doi:10.1186/s12864-017-3987-9)

26. Valladão R, Neto OBS, de Oliveira Gonzaga M, Pimenta DC, Lopes AR. 2023 Digestive enzymes and sphingomyelinase D in spiders without venom (Uloboridae). Sci. Rep. 13, 2661. (doi:10.1038/s41598-023-29828-x)

27. Zhang Y et al. 2024 A trade-off in evolution: the adaptive landscape of spiders without venom glands. GigaScience 13, giae048. (doi:10.1093/gigascience/giae048)

28. Kuhn-Nentwig L, Langenegger N, Heller M, Koua D, Nentwig W. 2019 The dual prey-inactivation strategy of spiders—in-depth venomic analysis of Cupiennius salei. Toxins 11, 167. (doi:10.3390/toxins11030167)

29. Abramson J et al. 2024 Accurate structure prediction of biomolecular interactions with AlphaFold 3. Nature 630, 493–500. (doi:10.1038/s41586-024-07487-w)

30. Dresler J, Avella I, Damm M, Dersch L, Krämer J, Vilcinskas A, Lüddecke T. 2024 A roadmap to the enzymes from spider venom: biochemical ecology, molecular diversity, and value for the bioeconomy. Front. Arachn. Sci. 3. (doi:10.3389/frchs.2024.1445500)

31. Hu H, Bandyopadhyay PK, Olivera BM, Yandell M. 2011 Characterization of the Conus bullatus genome and its venom-duct transcriptome. BMC Genomics 12, 60. (doi:10.1186/1471-2164-12-60)

32. Abalde S, Tenorio MJ, Afonso CML, Zardoya R. 2018 Conotoxin diversity in Chelyconus ermineus (Born, 1778) and the convergent origin of piscivory in the Atlantic and Indo-Pacific cones. Genome Biol. Evol. 10, 2643–2662. (doi:10.1093/gbe/evy150)

33. Dresler J, Avella I, Damm M, Dersch L, Krämer J, Vilcinskas A, Lüddecke T. 2024 A roadmap to the enzymes from spider venom: biochemical ecology, molecular diversity, and value for the bioeconomy. Front. Arachn. Sci. 3. (doi:10.3389/frchs.2024.1445500)

34. Zhu S, Peigneur S, Gao B, Luo L, Jin D, Zhao Y, Tytgat J. 2011 Molecular diversity and functional evolution of scorpion potassium channel toxins. Mol. Cell. Proteomics 10, S1– S11. (doi:10.1074/mcp.M110.002832)

35. Meng L et al. 2020 Ion channel modulation by scorpion hemolymph and its defensin ingredients highlights origin of neurotoxins in telson formed in Paleozoic scorpions. Int. J. Biol. Macromol. 148, 351–363. (doi:10.1016/j.ijbiomac.2020.01.133)

36. Zhu S, Peigneur S, Gao B, Umetsu Y, Ohki S, Tytgat J. 2014 Experimental conversion of a defensin into a neurotoxin: implications for origin of toxic function. Mol. Biol. Evol. 31, 546–559. (doi:10.1093/molbev/msu038)

37. Karnovsky MJ. 1965 A formaldehyde glutaraldehyde fixative of high osmolality for use in electron microscopy. J Cell Biol 27, 1A–149A.

38. Andrews S. 2010 FastQC: A quality control tool for high throughput sequence data. See https://www.bioinformatics.babraham.ac.uk/projects/fastqc/ (accessed on 4 June 2021).

39. Chen S, Zhou Y, Chen Y, Gu J. 2018 fastp: an ultra-fast all-in-one FASTQ preprocessor. Bioinformatics 34, i884–i890. (doi:10.1093/bioinformatics/bty560)

40. Bushmanova E, Antipov D, Lapidus A, Prjibelski AD. 2019 rnaSPAdes: a de novo transcriptome assembler and its application to RNA-Seq data. GigaScience 8, giz100. (doi:10.1093/gigascience/giz100)

41. Signal B, Kahlke T. 2021 Borf: Improved ORF prediction in de-novo assembled transcriptome annotation. BioRxiv, 2021.04.12.439551. (doi:10.1101/2021.04.12.439551)

42. Camacho C, Coulouris G, Avagyan V, Ma N, Papadopoulos J, Bealer K, Madden TL. 2009 BLAST+: architecture and applications. BMC Bioinformatics 10, 421. (doi:10.1186/1471-2105-10-421)

43. The UniProt Consortium. 2023 UniProt: the universal protein knowledgebase in 2023. Nucleic Acids Res. 51, D523–D531. (doi:10.1093/nar/gkac1052)

44. Pineda SS et al. 2018 ArachnoServer 3.0: an online resource for automated discovery, analysis and annotation of spider toxins. Bioinforma. Oxf. Engl. 34, 1074–1076. (doi:10.1093/bioinformatics/btx661)

45. Quevillon E, Silventoinen V, Pillai S, Harte N, Mulder N, Apweiler R, Lopez R. 2005 InterProScan: protein domains identifier. Nucleic Acids Res. 33, W116–W120. (doi:10.1093/nar/gki442)

46. Teufel F et al. 2022 SignalP 6.0 predicts all five types of signal peptides using protein language models. Nat. Biotechnol. 40, 1023–1025. (doi:10.1038/s41587-021-01156-3)

47. Miller J, Zimin AV, Gordus A. 2023 Chromosome-level genome and the identification of sex chromosomes in Uloborus diversus. GigaScience 12, giad002. (doi:10.1093/gigascience/giad002)

48. Postic G, Gracy J, Périn C, Chiche L, Gelly J-C. 2018 KNOTTIN: the database of inhibitor cystine knot scaffold after 10 years, toward a systematic structure modeling. Nucleic Acids Res. 46, D454–D458. (doi:10.1093/nar/gkx1084)

49. R Core Team. 2019 R: A language and environment for statistical computing. R Foundation for Statistical Computing, Vienna, Austria. URL http://www.R-project.org/. (doi:URL http://www.R-project.org/)

50. Gouy M, Guindon S, Gascuel O. 2010 SeaView Version 4: A Multiplatform graphical user interface for sequence alignment and phylogenetic tree building. Mol. Biol. Evol. 27, 221–224. (doi:10.1093/molbev/msp259)

51. Edgar RC. 2004 MUSCLE: multiple sequence alignment with high accuracy and high throughput. Nucleic Acids Res. 32, 1792–1797. (doi:10.1093/nar/gkh340)

52. Hassan A, Blakeley G, McGregor AP, Zancolli G. 2024 Venom gland organogenesis in the common house spider. Sci. Rep. 14, 15379. (doi:10.1038/s41598-024-65336-2)

53. Dobin A, Davis CA, Schlesinger F, Drenkow J, Zaleski C, Jha S, Batut P, Chaisson M, Gingeras TR. 2013 STAR: Ultrafast universal RNA-seq aligner. Bioinformatics 29, 15–21. (doi:10.1093/bioinformatics/bts635)

54. Soneson C, Love MI, Robinson MD. 2015 Differential analyses for RNA-seq: transcript-level estimates improve gene-level inferences. F1000Research 4, 1521. (doi:10.12688/f1000research.7563.1)

55. Fischer ML et al. 2024 Divergent venom effectors correlate with ecological niche in neuropteran predators. Commun. Biol. 7, 981. (doi:10.1038/s42003-024-06666-9)

56. Peigneur S et al. 2021 Small cyclic sodium channel inhibitors. Biochem. Pharmacol. 183, 114291. (doi:10.1016/j.bcp.2020.114291)

