## Supplementary for "Beyond venomous fangs: Uloboridae spiders have lost their venom but not their toxicity"

**Table S1**: RNA-seq library summary statistics. UC1-3: chelicerae; UP1-3: prosoma; UG: gonads; UM1-3: midgut glad; US1-3: silk glands.

| Libraries | Total raw reads | High quality reads |
| --- | --- | --- |
| UC1 | 58'494'780 | 48'903'542 |
| UC2 | 53'421'906 | 38'774'986 |
| UC3 | 54'406'220 | 38'713'640 |
| UP1 | 55'299'872 | 46'506'656 |
| UP2 | 75'303'620 | 67'291'910 |
| UP3 | 98'080'052 | 74'005'606 |
| UG1 | 107'756'820 | 96'386'224 |
| UM1 | 96'262'266 | 85'748'874 |
| UM2 | 90'920'880 | 83'279'616 |
| UM3 | 95'046'808 | 86'684'478 |
| US1 | 53'881'834 | 44'579'252 |
| US2 | 53'881'834 | 48'642'782 |
| US3 | 53'941'648 | 49'819'522 |

**Table S2**: *De novo* transcriptome assembly summary statistics. ORF: open reading frame.

|  | # Transcripts | |
| --- | --- | --- |
|  | **Complete ORF** | **Incomplete ORF** |
| Assembled sequences | 302'504 | |
| Transcripts with ORF | 159'668 | 66'125 |
| Annotated sequences | 33'129 | 260 |
| Clustered sequences* | 20'050 | 193 |

* After grouping sequences with 99% identity.

**Table S3**: List of the 11 spider genomes used for the assembly annotation.

| Assembly accession | Species | Total sequence length | Assembly level | Assembly submission date |
| --- | --- | --- | --- | --- |
| GCA_013235015.1 | *Araneus ventricosus* | 3'656'621'265 | Scaffold | 2019-08-02 |
| GCA_015342795.1 | *Argiope bruennichi* | 1'670'285'661 | Chromosome | 2020-11-16 |
| GCA_021605075.1 | *Caerostris darwini* | 1'501'919'382 | Scaffold | 2021-11-19 |
| GCA_021605095.1 | *Caerostris extrusa* | 1'420'656'204 | Scaffold | 2021-11-19 |
| GCA_019974015.1 | *Nephila pilipes* | 2'694'500'076 | Scaffold | 2021-07-23 |
| GCA_019343175.1 | *Oedothorax gibbosus* | 821'427'276 | Chromosome | 2021-08-05 |
| GCA_000365465.3 | *Parasteatoda tepidariorum* | 1'228'972'128 | Scaffold | 2019-06-14 |
| GCA_010614865.2 | *Stegodyphus dumicola* | 2'551'176'228 | Scaffold | 2020-02-14 |
| GCA_019973975.1 | *Trichonephila clavata* | 2'497'895'991 | Scaffold | 2021-07-23 |
| GCA_019973935.1 | *Trichonephila clavipes* | 2'874'350'602 | Scaffold | 2021-07-23 |
| GCA_019973955.1 | *Trichonephila inaurata madagascariensis* | 2'507'041'000 | Scaffold | 2021-07-22 |

**Table S4:** List of fastq files used to verify and quantify defensin in the genome of *Octonoba sinensis*.

| **SRA** | **Species** | **tissue** |
| --- | --- | --- |
| SRR26131816 | *Octonoba sinensis* | Brain |
| SRR26148751 | *Octonoba sinensis* | Abdomen |
| SRR26148754 | *Octonoba sinensis* | Brain |
| SRR26148755 | *Octonoba sinensis* | Brain |
| SRR26148758 | *Octonoba sinensis* | Chelicera |
| SRR26148759 | *Octonoba sinensis* | Chelicera |
| SRR26148770 | *Octonoba sinensis* | Silk gland |
| SRR26148771 | *Octonoba sinensis* | Silk gland |
| SRR26148772 | *Octonoba sinensis* | Silk gland |
| SRR26148773 | *Octonoba sinensis* | Gut |
| SRR26148774 | *Octonoba sinensis* | Gut |
| SRR26148775 | *Octonoba sinensis* | Gut |
| SRR26148776 | *Octonoba sinensis* | Abdomen |
| SRR26148777 | *Octonoba sinensis* | Abdomen |

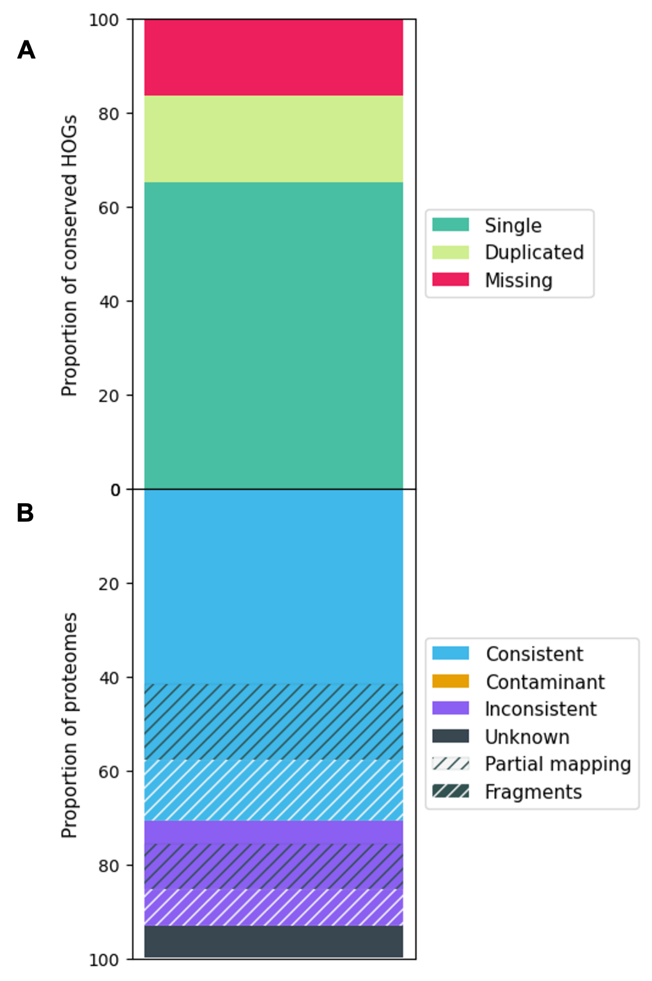

**Fig. S1 OMArk completeness evaluation of *U. plumipes* transcriptome assembly**. **A**. Proportion of conserved Hierarchical Ortholog groups (HOGs) using Arthropoda as ancestral clade which contains 3’589 conserved HOGs. Results on conserved HOGs: Single: 2’335 (65.06%); Duplicated: 662 (18.45%); Duplicated, Unexpected: 639 (17.80%); Duplicated, Expected: 23 (0.64%); Missing: 592 (16.49%). **B**. Proportion of proteins in the transcriptome with a consistent lineage placement: Total consistent, 14’328 (70.78%); Consistent, partial hits, 3’290 (16.25%); Consistent, fragmented: 2’617 (12.93%). Inconsistent lineage placements: Total inconsistent, 4’512 (22.29%); Inconsistent, partial hits: 1’948 (9.62%); Inconsistent, fragmented: 1’547 (7.64%). Total unknown: 1’403 (6.93%).

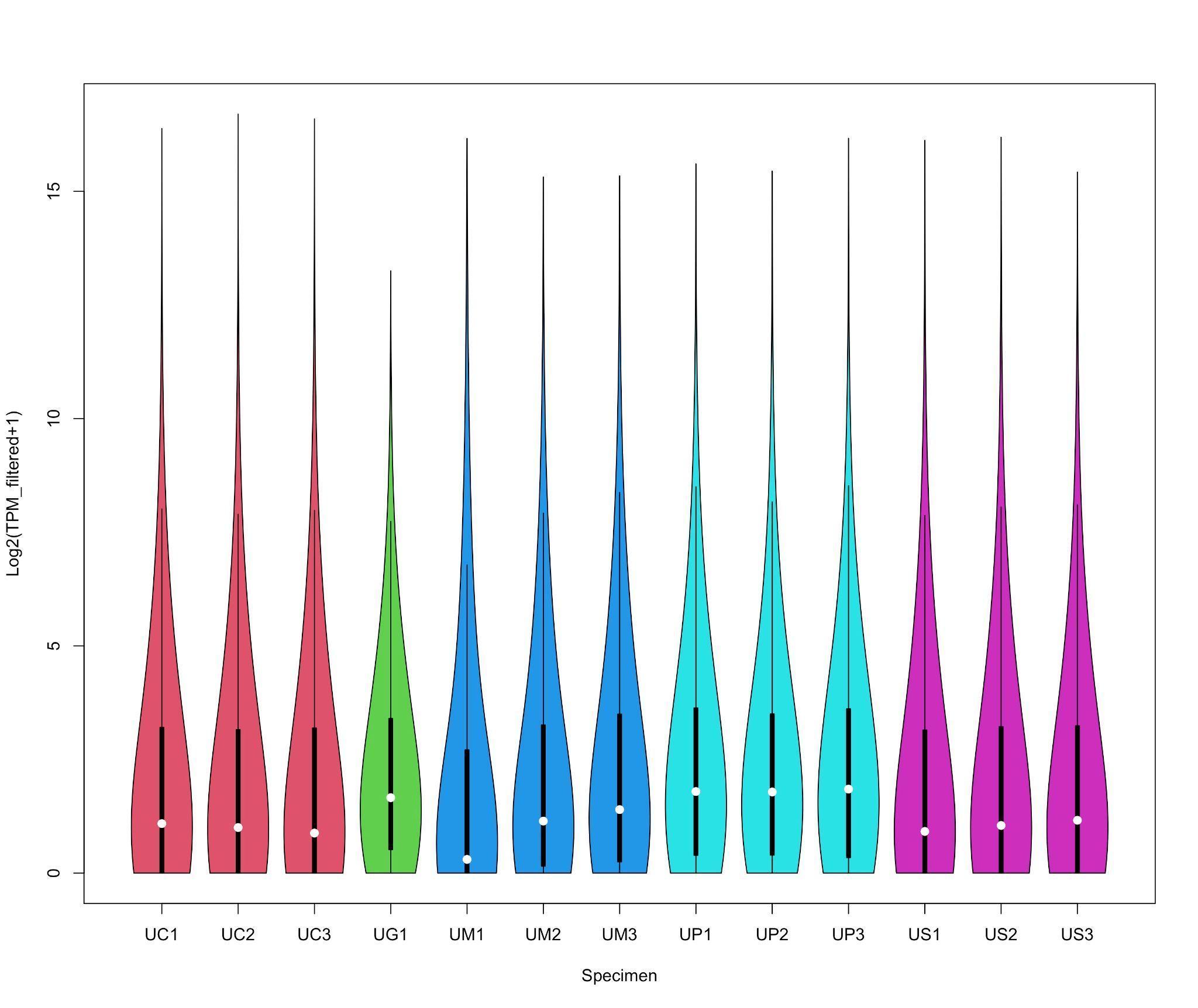

**Fig. S2 Violine plot of library transcripts per million (TPMs) data.** UC1, UC2, UC3: chelicerae; UG1: gonad; UM1, UM2, UM3: midgut gland; UP1, UP2, UP3: prosoma; US1, US2, US3: silk glands.

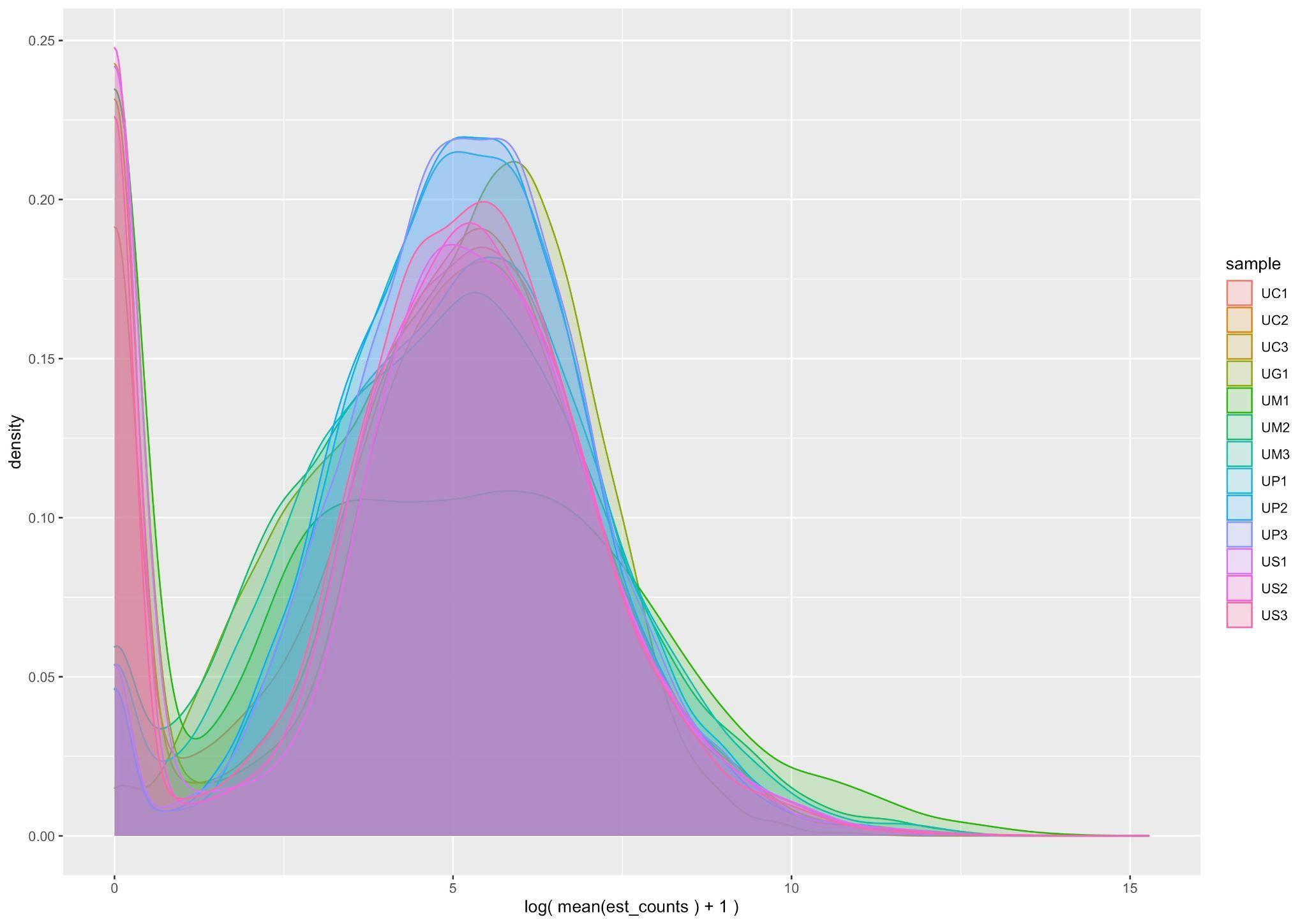

**Fig. S3 Read count density of libraries.** UC1, UC2, UC3: chelicerae; UG1: gonad; UM1, UM2, UM3: midgut gland; UP1, UP2, UP3: prosoma; US1, US2, US3: silk glands.

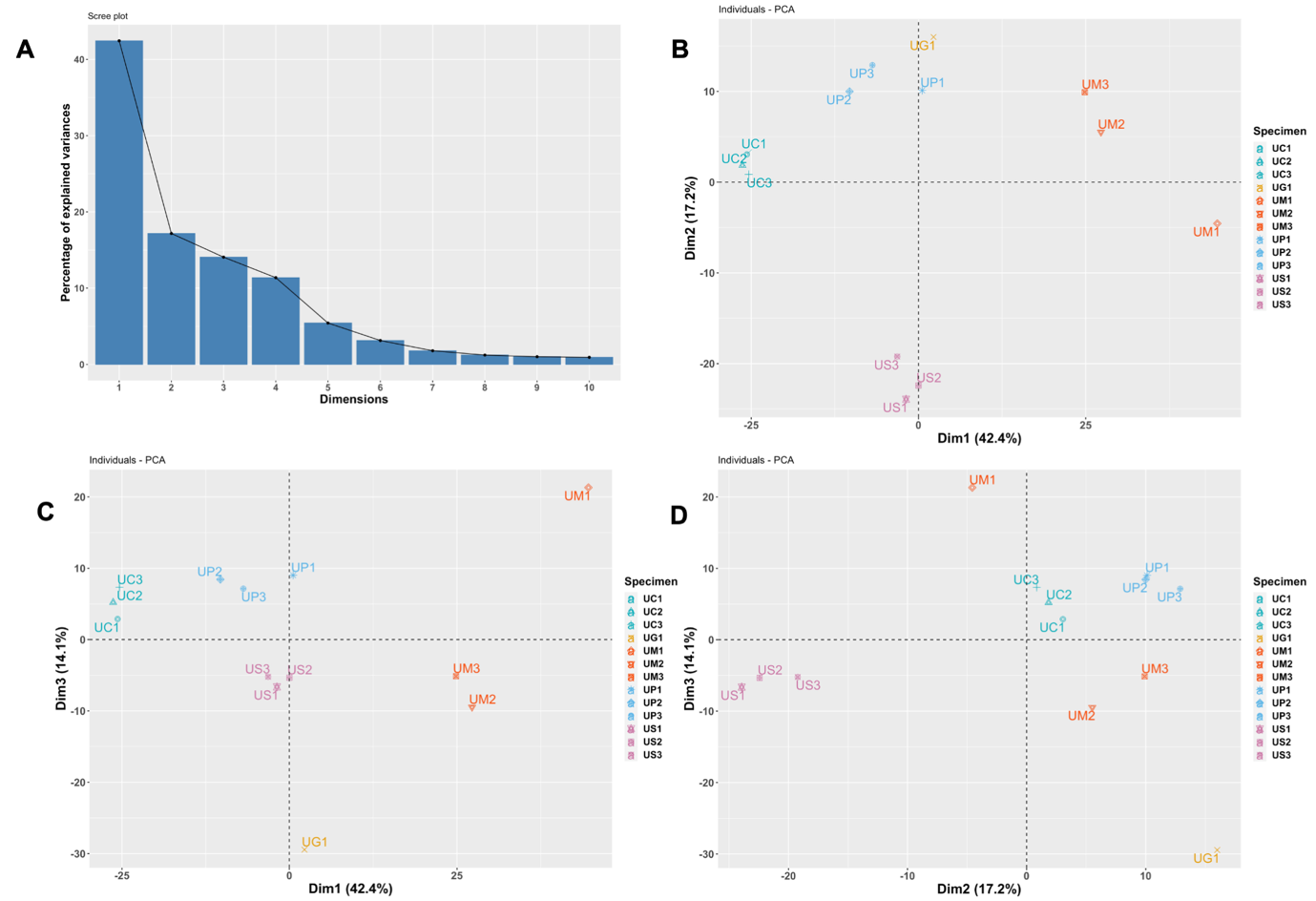

**Fig. S4 Principal component analysis using the top 1’000 most variable transcripts. A.** Scree plot of the components’ explained variance. **B, C, D**. PCA plot with the first two component (**B**), the first and third (**C**), and second and third components (**D**). UC1, UC2, UC3: chelicerae; UG1: gonad; UM1, UM2, UM3: midgut gland; UP1, UP2, UP3: prosoma; US1, US2, US3: silk glands.

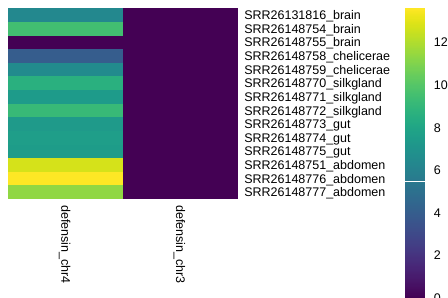

**Fig. S5. Expression levels of the newly annotated defensin genes in the *Octonoba sinensis* genome.** Expression levels as log_2_ (TPM).

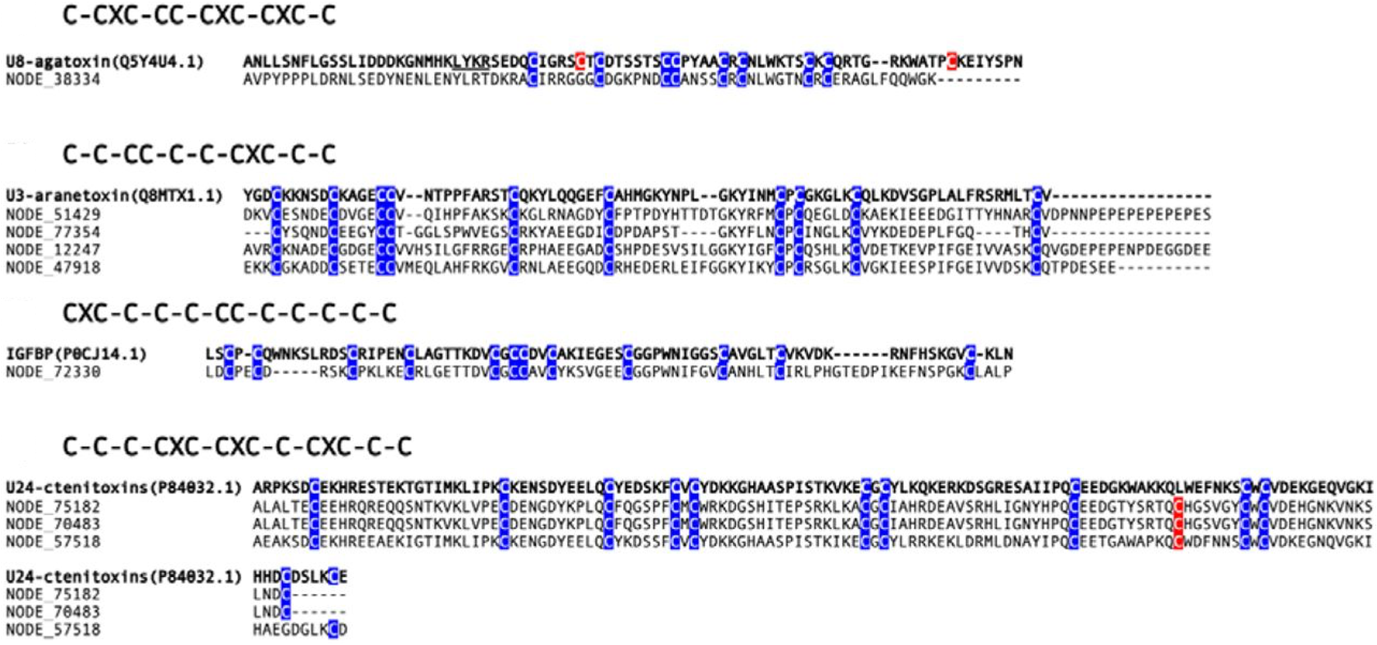

**Fig. S6. Alignment of predicted neurotoxins from the *U. plumipes* transcriptome.** The cysteine frame is reported.
